# Forecasting Dengue Fever in Brazil Using Multimodal Data, Including Climate Data

**DOI:** 10.1101/2024.03.12.584724

**Authors:** David C. Danko, Charles Vörösmarty, Anthony Cak, Fabio Corsi, Suraj Patel, John Papciak, Heather Wells, Anthony Albanese, Christopher E. Mason, Rafael Maciel-de-Freitas, Dorottya Nagy-Szakal, Niamh B. O’Hara

## Abstract

Dengue fever is a major tropical disease transmitted by *Aedes* mosquitoes. Dengue affects more than 120 countries with highly variable year to year infection rates. Despite high variability, dengue has a clear relationship to climate factors and human demography. Global trends to higher temperatures and greater disorderly urban development are increasing the scale and scope of dengue risk. Dengue has complex human immunity with 4 known serotypes that make multiple infections possible. Accurate forecasting of dengue fever would allow for appropriate interventions and improved public health outcomes. We demonstrate, *GeoSeeq Dengue*, a forecasting model for dengue fever in Brazil. GeoSeeq Dengue predicts dengue outbreaks monthly in 5,570 Brazilian municipalities at 1, 3, and 6 months ahead of the outbreak. Model accuracy compares favorably to a historical baseline model, making it a promising model for informing public health response. We evaluate how different types of input variables effect model accuracy and explore how this model could be adapted to other countries. This model could inform public health responses to dengue including targeting vector control programs, public health education messaging, and the newly launched dengue vaccine rollout.

## Introduction

Dengue fever is a major tropical disease transmitted principally by mosquitoes. Dengue has a burden of roughly 1,600 disability adjusted life years per million people with high year to year variability (Guzman 2010). The range of dengue is largely concomitant with the range of its mosquito vector species. Climate change is expanding that range into previously inhospitable regions. As such, dengue cases have been expanding and rising in recent years, particularly in Brazil (Barcellos 2024).

In Brazil, dengue is principally transmitted by *Aedes aegypti* and to a lesser extent *Aedes albopictus*. Both species are principally crepuscular and survive well in human built environments, though *aegypti* is more common in urban areas and *albopictus* in exurban settings.

Brazil is divided into 27 federal states comprising 5,570 municipios responsible for aspects of public health. Municipios cover the entire country (except for a federal district) and vary in climate, geographic size, and population. Brazil tracks dengue case rates at the municipio level, providing a long dense time series of case data, which is useful for predictive model training. The peak season for dengue in Brazil begins in January and lasts typically until June, though some municipios regularly experience peaks at other times. Until relatively recently the southern part of Brazil had few *Aedes* mosquitoes and little to no dengue.

Historically public health strategies to mitigate dengue have focused on mosquito spraying programs, mosquito net advisories, and increased care resources for affected areas. Recently, effective vaccines for dengue have become available, but will require a prolonged rollout to reach the large affected populations. Infection of *Aedes* mosquitoes with *Wolbachia* has also been shown to reduce carrying rates of dengue (and other arboviruses) and this has been proposed as a public health mitigation strategy.

One of the key challenges to any public health effort focusing on dengue is the high variability of the disease. Owing to complex human immunity, climate factors, human mobility trends, and random variability, the rate of dengue can vary widely both spatially and temporally. Effective forecasting for dengue would make allocation of public health resources more optimal and would lead to a reduced disease burden. Different types of intervention may require different forecasting time scales. To be effective it must also be possible to integrate a forecast into existing public health frameworks and to communicate results to decision makers.

There have been several efforts to forecast dengue in Brazil and elsewhere including: Sebastianelli 2024 which forecast dengue rates one month ahead at the state level, Kuo 2014 which forecasts dengue in Columbia from satellite imagery, Roster 2022 which predicted case rates in certain Brazilian cities 1-month ahead, Leandro 2022 which investigated the role of mosquitos in transmission in Foz do Iguaçu, and Lowe 2014 which predicted rates around the 2014 World Cup.

In *GeoSeeq Dengue* we have built a set of predictive models that includes predictions at multiple time points (1, 3, and 6 months) and at a more fine-grained scale, at the level of the municipio.

## Results

We developed a number of predictive models for dengue outbreaks in Brazil. These models are trained to predict if a given municipio would predict an outbreak in a particular month which we defined as a case rate greater than 350 cases per 100k people. Models were trained using data from 5,570 municipios from January 2001 through December 2016. Models were evaluated on the same 5,570 municipios with data from January 2017 through December 2019. Models were built to forecast 1-month, 3-months, and 6-months ahead. Longer forecasts had slightly less training and test data available to them due to the necessary lead data.

### Establishing a Baseline

We developed two non-random baseline models to provide useful comparisons to our predictive models. Dengue has strong historical and geographic trends-to be useful a predictive model should outperform simple predictors.

First, we developed a model that predicts that a municipio will continue to have an outbreak if it already is having one and will not have an outbreak if it is not. We call this the Same as Previous Month (**SAPM**) model. This model uses data from one month prior to the target month, by construction this means SAPM has data not available to the 3-month and 6-month forecasts. Performance of SAPM is reasonable with an F1 score of 58.8%.

Second, we developed a model based on the Historical Mean Case Rate (**HMCR**) of a municipio. We took the average case rate in each municipio for each month in our training period. We then applied a threshold to the case rate, if a municipio’s mean case rate was above this threshold that month was considered to be an outbreak in that municipio. HMCR performed substantially worse than SAPM with a best F1 score of 21.6%. We note that this model produces the same result regardless of how far ahead of time the predictions are made.

### Past Year Case Rates

We developed a model that predicts dengue outbreaks in a municipio based on the case rates in that municipio from the 12 months prior to the prediction as well as the month being predicted (e.g. March). We term this the Past Year Case Rate (**PYCR**) model. Dengue outbreaks have a characteristic shape with low case rates leading exponentially to a peak. This peak can sustain for 1-3 months then will drop off exponentially. Recent case rate data could be used to identify municipios in the early stages of outbreaks or to identify municipios in the intermediate or end stages. Since the model also has access to the target month it can learn seasonal patterns as well.

We trained and tested versions of the model predicting 1, 3, and 6 months ahead. For example, the 3-month model would predict an outbreak in March 2017 in a single municipio given the case rates Jan-Dec 2016 from only that municipio. This model shows good performance 1 month ahead with performance degrading substantially as the timeframe increases. At 1 month PYCR has better performance than the SAPM. Three and six months ahead performance exceeds the HMCR baseline model but is worse than the SAPM baseline (which uses data inaccessible to these forecasts).

We use our PYCR model as the basis for further modeling. By supplementing the PYCR model with other feature sets we are able to improve overall performance.

**Table 1:** Performance of dengue forecasting models. Gives the precision, recall, best F1 score, and the Area Under the Precision Recall Curve (AUG) for each model. Note that this AUG differs slightly from AUG of the ROG curve, the metric typically used in balanced classification problems.

| Model | Forecast Time | AUC (Pr/Re) | F1 | Precision | Recall |
| --- | --- | --- | --- | --- | --- |
| HMCR (Baseline) | Infinite | 13.4% | 21.6% | 16.0% | 33.1% |
| SAPM (Baseline) | Months Ahead: 1 | 44.7% | 58.8% | 56.7% | 61.0% |
| Geophysical | Months Ahead: 1 | **72.6%** | **67.0%** | 66.9% | 67.1% |
| Geospatial Variables | Months Ahead: 1 | 72.0% | 66.9% | 67.7% | 66.1% |
| Historical Rate | Months Ahead: 1 | 65.9% | 62.3% | 62.3% | 62.4% |
| Mosquito Activity | Months Ahead: 1 | 72.4% | 67.2% | 66.1% | 68.4% |
| PYCR | Months Ahead: 1 | 70.6% | 66.2% | 65.7% | 66.6% |
| Geophysical | Months Ahead: 3 | **34.0%** | 39.0% | 36.3% | 42.2% |
| Geospatial Variables | Months Ahead: 3 | 32.4% | 38.5% | 34.8% | 43.1% |
| Historical Rate | Months Ahead: 3 | 29.5% | 34.1% | 29.3% | 40.7% |
| Mosquito Activity | Months Ahead: 3 | 33.5% | **39.1%** | 34.0% | 46.0% |
| PYCR | Months Ahead: 3 | 26.0% | 33.8% | 29.8% | 39.2% |
| Geophysical | Months Ahead: 6 | **26.0%** | **33.1%** | 26.0% | 45.5% |
| Geospatial Variables | Months Ahead: 6 | 17.0% | 23.8% | 17.8% | 35.9% |
| Historical Rate | Months Ahead: 6 | 24.4% | 32.2% | 26.3% | 41.6% |
| Mosquito Activity | Months Ahead: 6 | 23.4% | **33.1%** | 26.1% | 45.1% |
| PYCR | Months Ahead: 6 | 19.6% | 29.1% | 21.8% | 43.5% |

### Geospatial Data

We extended our PYCR model by including case rate data from other parts of Brazil. Dengue has regional trends that relate outbreaks in nearby and distant municipios. By providing a machine learning model with regional case rate information it can learn these geospatial trends. We implement this by dividing Brazil’s municipios into 16 clusters. We aggregate the average case rate in each cluster for each of the three months prior to the prediction. We train the model to predict an outbreak in a municipio from the past 12 months of case rates in that municipio, the month being predicted, three months of aggregated case rates from each cluster, and an index giving cluster membership of the target municipio.

Including geospatial case rate data provides a slight performance improvement over the 1-month PYCR model and a substantial improvement over the 3-month PYCR model. However, performance is actually degraded compared to the 6-month PYCR model despite providing strictly more information (likely a consequence of chosen model architecture). However, 6-month performance still exceeded the HMCR baseline.

### Geophysical Data

We extended our PYCR model by including geophysical variables from the target municipios. The geophysical variables used were closely related to temperature and rainfall. Variables were aggregated from raw measurements for each municipio in the time period leading up to the prediction. We note that we only used actual geophysical measurements not forecasts of the geophysical variables (though good forecasts for geophysical variables are often available).

Including geophysical data with our PYCR model provided a slight performance improvement over the 1-month PYCR model and a substantial improvement over the 3-month and 6-month PYCR model. Notably, performance of the 6-month model exceeded the HMCR baseline model.

### Mosquito Model

We extended our PYCR model by including a model of *Aedes* mosquito activity in target municipios. This model estimates the activity of *Aedes aegypti* and *Aedes albopictus* mosquitoes in a municipio in the time period before the prediction is made. The mosquito model is itself based on geophysical data, and population data, and is conditioned on mosquito observation data.

Including the mosquito model with our PYCR model provided a slight performance improvement over the 1-month PYCR model and a substantial improvement over the 3-month and 6-month PYCR model. Performance was similar to the performance achieved by using geophysical variables directly. Notably performance was improved when we included estimates of *albopictus* and *aegypti* as separate variables rather than one aggregated number.

### Historical Rates

We extended our PYCR model by including historical case rate data for the target municipios. This is essentially the same information used to build our HMCR baseline model. Performance 1-month ahead was slightly degraded despite being strictly more information. Performance 3-months out was slightly improved and performance 6-months ahead was substantially better than both PYCR and HMCR.

### Model Calibration

We tested our models to determine if they matched overall dengue trends in Brazil. We compared the predicted outbreak rate of our PYCR+Geophysical models to the actual outbreak rate for the three years in our test period (fig. 2A). All three prediction periods correctly anticipated the major 2019 outbreak relative to the more mild 2017 and 2018 dengue seasons. The 3-month model also correctly distinguished the 2017 and 2018 outbreaks.

**Figure 1:**
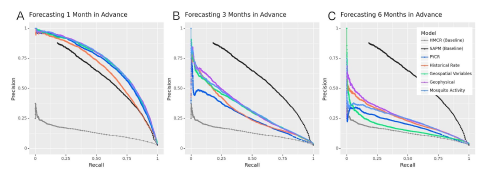
Effect of different variables on forecasting performance. Each panel shows precision recall curves for two baseline models (HMCR and SAPM), the core PYCR model alone, and the PYCR model augmented with four different sets of variables. Panel A shows model performance forecasting 1 month ahead, panel B 3 months, and panel C 6 months. We note that the SAPM baseline uses data 1 month prior to the forecasted month even for panels B & C. Color key is the same for all panels.

**Figure 2:**
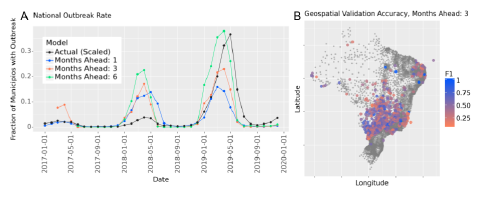
A) Fraction of municipios experiencing outbreaks over time. Predicted rates scale with actual rate. Actual rate shown as 2x for clarify. B) Geospatial accuracy per municipio. Points show F1 statistic at the geographic center of each municipio. Grey points are municipios that did not experienece an outbreak in the validation period.

We also tested the spatial distribution of accuracy for our PYCR+Geophysical models. We compared the F1 score for each municipio to the geographic center of each municipio (fig. 2B). Broadly, accuracy scores are evenly distributed across the country but not enough municipios outside of the central region had outbreaks during the validation period for a comprehensive analysis. The spatial distribution of scores was broadly similar at all three forecasting timeframes.

### Case Rate Free Models

Brazil has accurate nationwide records of dengue case rates going back for several decades. These records are both a valuable resource to build predictive models and a useful input to those models. However, in many other regions accurate case rate data may be inaccessible or may only go back a comparatively brief period.

In these regions models based on recent case rates (such as our PYCR model) would not be beneficial. To address this issue we developed a model based on geophysical variables and estimated mosquito activity but without any case rate data (case rate data was still used to train and evaluate the model, just not as input to the predictor).

This model performed similarly 1-month, 3-months, and 6-months out with the worst performance 1-month out (fig. 3). Overall performance was slightly better than HMCR 3 and 6 months ahead but 1 month ahead was worse. We note that in the setting where case rate data is not available at all the HMCR model would no longer be a reasonable baseline as it relies itself on case rate data.

**Figure 3:**
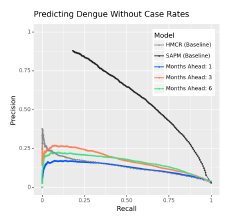
Models developed without case rate data can still provide a useful signal. Geophysical variables alone allow for better prediction than historical averages.

## Methods

### Data Access

Climate and geophysical data were accessed through NOAA and Google Earth Engine. Case rate data was accessed through SINAN. Population data was accessed through IBGE. Mosquito presence data was accessed from inaturalist and GBIF.

### Modeling Mosquito Activity

We reimplemented the model described by Musah 2023 to model *Aedes* mosquito activity in Brazil.

### Machine Learning Approach

All forecasting models in this paper (except for the baseline models) are random forest models as implemented in sklearn with 1,000 estimators and all other parameters at defaults. No attempt to tune model type or hyperparameters was made. Models were trained on data from all municipios. No finetuning to specific municipios or regions was performed.

## Discussion

We have developed GeoSeeq Dengue, a set of machine learning predictive models for dengue fever in Brazil. These models forecast outbreaks at the municipio level 1, 3, and 6 months in advance and outperform reasonable baseline models. We trained these models on data from 2001-2016 and validated them on data from 2017-2019. Models were, largely based on case rate data from the prior year supplemented with additional variables such as: mosquito activity, geophysical or climate variables, geospatial rates, and historical rates.

The forecast timeframe made a substantial and important difference in our model’s performance. Forecasting 1-month ahead was dominated by case rates from the prior year. While inclusion of other variables did improve accuracy, the scale of improvement was marginal. However, 3 and 6 months out the situation was quite different. In these scenarios recent case rates had much less predictive power and the inclusion of other variables improved accuracy much more substantially.

Of particular interest was the benefit of geophysical variables and models of mosquito activity to longer range forecasts. Climatic variables can be readily measured via satellite and our data suggests that even quite coarse application of these variables can lead to greatly improved outcomes. Interestingly, when these variables are compressed into a model of mosquito activity they retain most of their predictive power. Mosquito lifecycles are well understood by traditional biological approaches and this suggests that that knowledge can be used to effectively enhance larger scale predictive models.

Brazil has quite detailed case records compared to most other regions. These records are clearly valuable as a model input yet non-trivial forecasts can still be made without them. This is promising for two reasons. First, it suggests that it will be possible to build models of dengue in regions with less historical case information. Second, it suggests that it will be possible to build high resolution models of dengue that are finer than the resolution that records are collected at (municipio in Brazil).

Future work will include developing a more accurate optimized version of the models presented here, developing higher resolution models, and developing models that account for the complexities of human immunity with dengue serotypes.

## Data Availability

Data, models, and additional materials can be found on GeoSeeq at https://portal.geoseeq.com/sample-groups/880ce6b4-d09f-4a0c-b175-03d084c1c983

## References

Guzman, M. G., Halstead, S. B., Artsob, H., Buchy, P., Farrar, J., Gubler, D. J., Hunsperger, E., Kroeger, A., Margolis, H. S., Martínez, E., Nathan, M. B., Pelegrino, J. L., Simmons, C., Yoksan, S., & Peeling, R. W. (2010). Dengue: a continuing global threat. Nature reviews. Microbiology, 8(12 Suppl), S7–S16. 10.1038/nrmicro2460

Barcellos, C., Matos, V., Lana, R. M., & Lowe, R. (2024). Climate change, thermal anomalies, and the recent progression of dengue in Brazil. Scientific reports, 14(1), 5948. 10.1038/s41598-024-56044-y

Sebastianelli, A., Spiller, D., Carmo, R., Wheeler, J., Nowakowski, A., Jacobson, L. V., Kim, D., Barlevi, H., Cordero, Z. E. R., Colón-González, F. J., Lowe, R., Ullo, S. L., & Schneider, R. (2024). A reproducible ensemble machine learning approach to forecast dengue outbreaks. Scientific reports, 14(1), 3807. 10.1038/s41598-024-52796-9

Kuo, K.-T., Moukheiber, D., Ordonez, S. C., Restrepo, D., Paddo, A. R., Chen, T.-Y., … Celi, L. A. (2024). DengueNet: Dengue Prediction using Spatiotemporal Satellite Imagery for Resource-Limited Countries. arXiv [Cs.CV]. Retrieved from http://arxiv.org/abs/2401.11114

Roster, K., Connaughton, C., & Rodrigues, F. A. (2022). Machine-Learning-Based Forecasting of Dengue Fever in Brazilian Cities Using Epidemiologic and Meteorological Variables. American journal of epidemiology, 191(10), 1803–1812. 10.1093/aje/kwac090

Leandro, A. S., de Castro, W. A. C., Lopes, R. D., Delai, R. M., Villela, D. A. M., & de-Freitas, R. M. (2022). Citywide Integrated Aedes aegypti Mosquito Surveillance as Early Warning System for Arbovirus Transmission, Brazil. Emerging infectious diseases, 28(4), 701–706. 10.3201/eid2804.211547

Lowe, R., Coelho, C. A., Barcellos, C., Carvalho, M. S., Catão, R.deC., Coelho, G. E., Ramalho, W. M., Bailey, T. C., Stephenson, D. B., & Rodó, X. (2016). Evaluating probabilistic dengue risk forecasts from a prototype early warning system for Brazil. eLife, 5, e11285. 10.7554/eLife.11285

Musah, A., Browning, E., Aldosery, A., Valerio Graciano Borges, I., Ambrizzi, T., Tunali, M., … Kostkova, P. (2023). Coalescing disparate data sources for the geospatial prediction of mosquito abundance, using Brazil as a motivating case study. Frontiers in Tropical Diseases, 4. doi:10.3389/fitd.2023.1039735

